# Oligodendrocyte precursor cell death does not contribute to lifelong population homeostasis

**DOI:** 10.64898/2026.10.28.762467

**Authors:** Yasmine Kamen, Aine Scholand, Robert A. Hill

**Affiliations:** Department of Biological Sciences, Dartmouth College, Hanover, NH 03755, USA

## Abstract

Oligodendrocyte precursor cells (OPCs) proliferate and differentiate into myelinating oligodendrocytes throughout life, but it is unclear if their continuous self-renewal is balanced by cell death. We find that OPCs do not die during development or adulthood, nor during remyelination, acute systemic inflammation, or aging. These findings challenge the assumption that spontaneous OPC death is common and reveal that OPC homeostasis does not depend on programmed cell death.

## Main

Oligodendrocyte precursor cells (OPCs) tile the central nervous system^1,2^ and differentiate into myelinating oligodendrocytes throughout life^3^. Maintaining their population homeostasis through proliferation is therefore critical for lifelong myelination^4,5^, myelin repair^6^, and to support cognitive function^7^. Proliferation of neural stem cells or neural progenitors is thought to be balanced by programmed cell death^8^. However, it is unclear if OPC self-renewal is coupled with cell death. Like neurons^9^, microglia^10^, and astrocytes^11^, oligodendrocyte lineage cells are thought to be overproduced and undergo a large scale death event during early postnatal development^12^. Mature oligodendrocytes are long lived and only die in aging^4,5,13,14^. In contrast, differentiating oligodendrocytes regularly undergo programmed cell death^5,15–18^. The evidence supporting OPC death is more ambiguous. The elimination of the first wave of OPCs from the dorsal forebrain has been suggested to indicate OPC death but this was not formally tested^19–21^ and the disappearance of these cells is disputed^22,23^. Moreover, studies report either little to no dying OPCs^15,17,24–29^ or varying proportions of dying OPCs^2,5,30–32^. Assumptions that OPCs do or do not die are built into our current knowledge of OPC homeostasis^2,24,33^, differentiation dynamics^16,18,32^, and interactions with neurons^21^, and therefore inform our understanding of OPC and oligodendrocyte dynamics and function.

Thus, we investigated if OPCs spontaneously undergo programmed cell death. We first performed longitudinal intravital imaging of OPCs and their progeny in the somatosensory cortex of two-month-old *Cnp*-mEGFP : NG2-CreER™ : Ai9 mice. Daily imaging allowed fate mapping of tdTomato^+^ cells as they remained OPCs, differentiated into mEGFP^+^ oligodendrocytes, or died (Fig. 1a,b). To determine at what stage of maturation cell death occurs, we analyzed the soma area and circularity of tdTomato^+^ cells on the day before they became mEGFP^+^ oligodendrocytes or died, and compared these values to control cells that remained OPCs. Consistent with previous studies^15,26,34–37^, cells that would become mEGFP^+^ oligodendrocytes had a larger and more circular soma than cells that remained OPCs (Fig. 1c). Cells about to die were slightly larger and more circular than OPCs, but smaller and less circular than cells becoming mEGFP^+^ oligodendrocytes, and did not statistically differ from either group, suggesting that they were in an intermediate stage, and therefore, that differentiation had begun prior to cell death. This is consistent with previous work suggesting that mitochondria remodeling in dying oligodendrocyte lineage cells resembles the changes observed upon OPC differentiation^35^.

**Figure 1.**
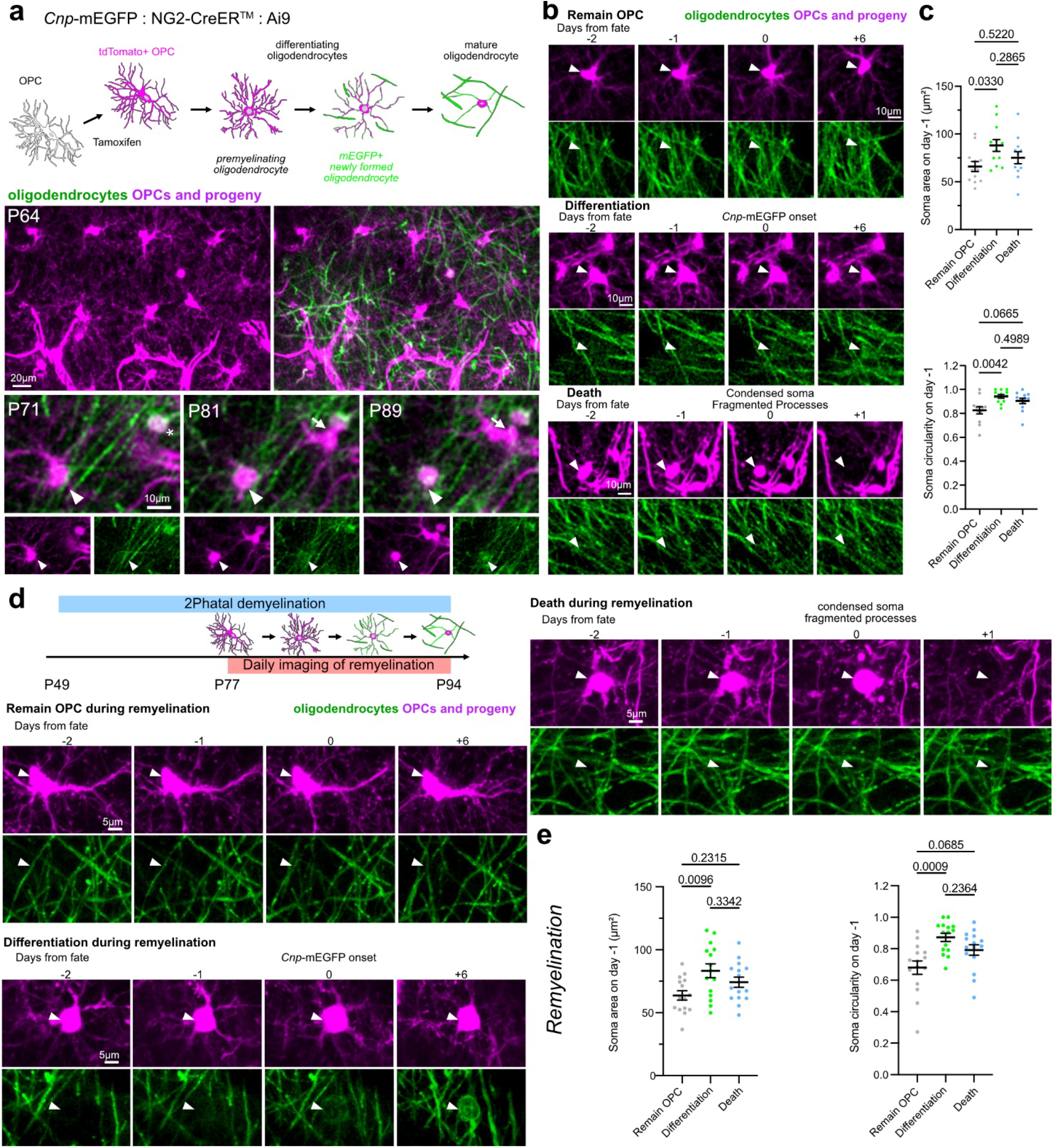
OPC differentiation precedes programmed cell death. **a** Schematic of the different maturation stages in the oligodendrocyte lineage. Tamoxifen injections in *Cnp*-mEGFP : NG2-CreER™ : Ai9 mice allow OPC and oligodendrocyte fate mapping, as shown in the example intravital images of oligodendrocytes (green) and OPCs and their progeny (magenta). The arrowhead indicates an OPC that differentiates and becomes mEGFP^+^. The arrow indicates an OPC that migrates into the region of interest. The asterisk indicates a mature oligodendrocyte. **b** Example of a cell that remains an OPC, differentiates to become mEGFP^+^, or dies. **c** Soma area and soma circularity on the day before cells became *Cnp*-mEGFP^+^ (Differentiation; green), died (Death; blue), or remained OPCs (grey) (n=12 cells from 4 mice, one-way ANOVA with Tukey’s multiple comparisons test). **d** Demyelination was induced with 2Phatal and daily imaging and fate mapping of OPCs were performed during remyelination. Examples of a cell that remained an OPC, differentiated, or died during remyelination^14^. **e** Soma area and soma circularity on the day before cells became *Cnp*-mEGFP^+^ (Differentiation; green), died (Death; blue), or remained OPCs (grey) during remyelination (n=15 cells from 2 mice, one-way ANOVA with Tukey’s multiple comparisons test). All data are shown as mean ± SEM.

OPC death is reported to be a feature of remyelination following ethidium bromide-induced demyelination^38^. We therefore wondered if OPC death was a general feature of remyelination. To address this, we reanalyzed the fate of tdTomato^+^ OPCs during remyelination in an intravital imaging dataset^14^ in which we used two-photon chemical apoptotic targeted ablation (2Phatal)^39^ to induce single oligodendrocyte death and demyelination in *Cnp*-mEGFP : NG2-CreER™ : Ai9 mice. We identified cells that remained OPCs, differentiated, or died in the 28 to 45 days following oligodendrocyte 2Phatal and measured their soma area and circularity (Fig. 1d,e). We found that cells about to die were larger and more circular than OPCs, but smaller and less circular than cells becoming mEGFP^+^ oligodendrocytes, and did not statistically differ from either group. These data suggest that during remyelination, as in homeostatic tissue, differentiation begins before programmed cell death occurs.

To refine our characterization of oligodendrocyte lineage stages undergoing spontaneous cell death, we turned to fixed tissue and used molecular markers to precisely define oligodendrocyte maturation. We first tested if we could detect PDGFRA^+^ OPCs with the apoptotic morphology we observed during intravital imaging (Fig. 2a; condensed soma, condensed nucleus, fragmented processes). We quantified the density of OPCs in the upper layers of the somatosensory cortex and corpus callosum between P4 and P32 and in one-year-old mice and classified each cell as having either a normal morphology or an apoptotic morphology. OPC density gradually decreased across development, as previously reported^21,40^ (Fig. 2b). However, OPC death did not underlie this decreased density, as only one out of 13,871 cortical OPCs (0.007%) and one out of 2,823 callosal OPCs (0.04%; Fig. 2a,b) displayed an apoptotic morphology. To ensure that we did not miss dying OPCs by analyzing morphology alone, we turned to cleaved caspase-3^+^ (CC3^+^) immunostaining. We focused on CC3 as differentiating oligodendrocytes regularly undergo CC3-mediated apoptosis^15,18,26^, and CC3^+^ OPCs have been reported^30,31^. There were numerous CC3^+^ cells in the somatosensory cortex of neonatal mice, but these were rarely OPCs (Fig. 2c,d; Extended Data Fig. 1). We found a total of three cortical CC3^+^ OPCs, representing 0.02% of OPCs counted between P4 and one year (Fig. 2c,d). Similarly, we only detected two callosal CC3^+^ OPCs, at P32 and one year. However, both cells were faintly BCAS1^+^, indicating that they may have started differentiating (Extended Data Fig. 2).

**Figure 2.**
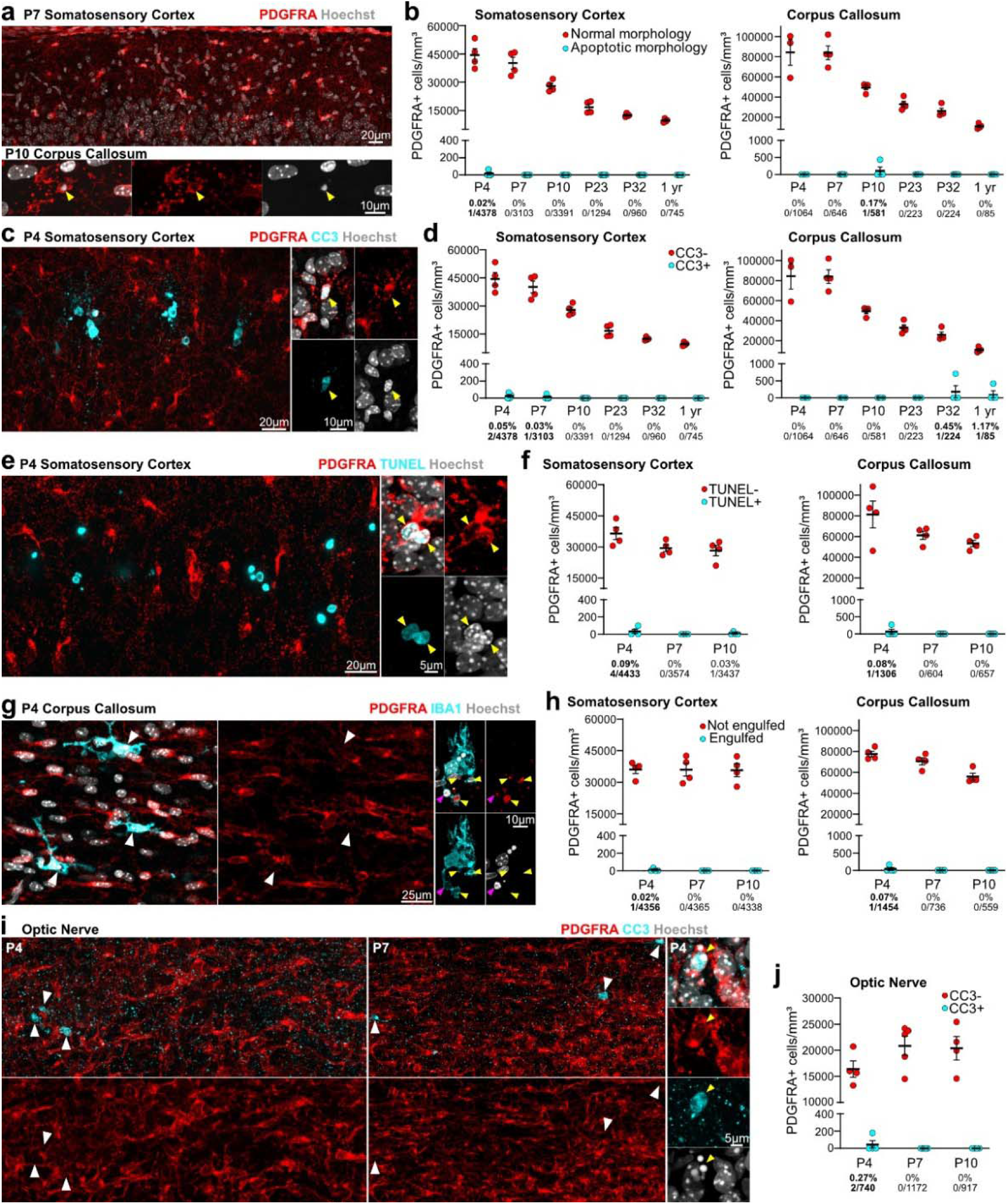
Dying OPCs are rarely detected. **a** PDGFRA (red) and nuclear dye (grey) staining allows the detection of OPCs. Most OPCs have a normal morphology (top), but a small number of OPCs appear apoptotic (bottom; yellow arrowhead; condensed soma, condensed and smooth nucleus, fragmented processes). **b** Density of PDGFRA^+^ OPCs with a normal (red) or apoptotic morphology (cyan) (n=3-4 mice). The numbers below the x axis indicate the proportion of PDGFRA^+^ cells with an apoptotic morphology/PDGFRA^+^ cells. **c** PDGFRA (red) and CC3 (cyan) rarely overlap (left), although the rare PDGFRA^+^CC3^+^ cell can be detected (right; yellow arrowhead). **d** Density of PDGFRA^+^CC3^-^ OPCs (red) and PDGFRA^+^CC3^+^ OPCs (cyan) (n=3-4 mice). The numbers below the x axis indicate the proportion of PDGFRA^+^CC3^+^ cells/PDGFRA^+^ cells. **e** TUNEL labeling (cyan) and PDGFRA staining (red) rarely overlap (left), but PDGFRA^+^TUNEL^+^ cells are occasionally detected (right; yellow arrowheads). **f** Density of PDGFRA^+^TUNEL^-^ OPCs (red) and PDGFRA^+^TUNEL^+^ OPCs (cyan) (n=4 mice). The numbers below the x axis indicate the proportion of PDGFRA^+^TUNEL^+^ cells/PDGFRA^+^ cells. **g** Left: Image of microglia (IBA^+^; cyan) engulfing nuclei (grey). These are highlighted with white arrowheads and do not overlap with PDGFRA (red). Right: rare example of microglia engulfing a PDGFRA^+^ OPC. The white arrowhead indicates the engulfed condensed OPC nucleus and the yellow arrowheads indicate the engulfed OPC apoptotic bodies. **h** Density of PDGFRA^+^ OPCs that are engulfed by microglia (cyan) or not engulfed by microglia (red) (n=4 mice). The numbers below the x axis indicate the proportion of engulfed PDGFRA^+^ cells/PDGFRA^+^ cells. **i** CC3 (cyan; indicated by white arrowheads) and PDGFRA (red) staining rarely overlap in the neonatal optic nerve (left), although rare examples of PDGFRA^+^CC3^+^ OPCs can be detected (right; yellow arrowhead). **j** Density of PDGFRA^+^CC3^-^ OPCs (red) and PDGFRA^+^CC3^+^ OPCs (cyan) (n=4-5 mice). The numbers below the x axis indicate the total proportion of PDGFRA^+^CC3^+^ cells/PDGFRA^+^ cells. All data are shown as mean ± SEM.

Unlike the lack of OPC death, both morphology analysis and CC3 labeling revealed a stable low density of dying differentiating oligodendrocytes (BCAS1^+^ cells) in the cortex between P4 and P32 and at one year, and in the corpus callosum between P4 and P10 (Extended Data Fig. 1a-d). This is consistent with low daily death rates we and others^5^ have observed with intravital imaging (0.18±0.04 death events/day in 4 two-month-old *Cnp*-mEGFP : NG2-CreER™ : Ai9 mice imaged over 21 days) but differs from reports of higher proportions of fragmented cortical differentiating oligodendrocytes during development^17^. These differences could be due to sampling differentiating oligodendrocytes at the leading edge of myelination, while we considered differentiating oligodendrocytes across all upper cortical layers.

Proportionally, there were 132-fold more CC3^+^ differentiating oligodendrocytes than CC3^+^ OPCS in the cortex between P4 and one year (differentiating oligodendrocytes: 2.64%, OPCs: 0.02%; p<1×10^-15^, χ^2^ test). Similarly, there were proportionally more CC3^+^ differentiating oligodendrocytes than CC3^+^ OPCs in the corpus callosum between P4 and P10 (differentiating oligodendrocytes: 1.09%, OPCs: 0%; p=2.2×10^-6^, χ^2^ test). We did not detect any CC3^+^BCAS1^-^CNPase^+^ cells (Extended Data Fig. 1e,f), consistent with previous work indicating that mature oligodendrocytes do not undergo spontaneous CC3-mediated apoptosis^5,14,26^.

It is possible that OPCs undergo a different form of cell death than CC3-mediated apoptosis. To expand the range of cell death mechanisms, we turned to Terminal Deoxynucleotidyl Transferase-dUTP Nick End Labeling (TUNEL). Although TUNEL is typically associated with apoptosis, we found that TUNEL and CC3 labeling did not always overlap in dying neurons (Extended Data Fig. 3). This is consistent with TUNEL labeling DNA strand breaks and therefore most forms of cell death involving DNA damage^41^. We focused on neonatal timepoints (P4-P10), as those were the timepoints in which we could detect the few dying OPCs, and like with CC3 staining, we found very few TUNEL^+^ OPCs in the neonatal cortex (four TUNEL^+^ OPCs out of 11,444 OPCs; 0.03%) or corpus callosum (one TUNEL^+^ OPC out of 2,567 OPCs; 0.04%; Fig. 2e,f). To exclude the possibility that OPCs spontaneously die without damaged DNA and through a CC3-independent mechanism, we assessed OPC engulfment by microglia^32^, as phagocytic clearance should occur independently of the death mechanism. As with the other death detection methods, we only found one cortical OPC (out of 13,059; 0.008%) and one callosal OPC (out of 2,749; 0.04%) engulfed by microglia, indicating that the vast majority of OPCs are not dying (Fig. 2g,h). Collectively, these data indicate that spontaneous programmed OPC death is extremely rare in the developing and adult somatosensory cortex and corpus callosum.

Extensive oligodendrocyte lineage cell death has been reported in the developing optic nerve^12^. We therefore tested if OPCs die in this white matter tract. As we detected similar numbers of dying OPCs in the cortex and corpus callosum with morphological analysis, CC3 staining, TUNEL labeling, and microglial engulfment, we examined OPC death in the optic nerve with CC3 staining only. CC3^+^ cells were distributed throughout the optic nerve and rarely overlapped with PDGFRA (Fig. 2i; Extended Data Fig. 1g,j). Between P4 and P10, we only detected two CC3^+^ OPCs out of 2,829 OPCs counted (0.07%; Fig. 2i,j). When examining cell death in oligodendrocyte lineage cells (SOX10^+^), we detected eight SOX10^+^CC3^+^ cells out of 6,609 SOX10^+^ cells (0.09%; Extended Data Fig. 1h,i). Thus, overall, we detected little OPC or oligodendrocyte lineage cell death in the developing optic nerve. This conflicts with the previous estimate, based on oligodendrocyte production estimates and increased cell numbers after growth factor addition, that 50% of oligodendrocytes die daily in the developing optic nerve^12^.

Microglia phagocytosis of cellular debris is efficient, and it is possible that most dying oligodendrocyte lineage cells are cleared too rapidly for immunohistochemical detection^12,33^. To address this possibility, we counted dying OPCs in the somatosensory cortex of P7 *Csf1r*^ΔFIRE^ mice. In these mice, the deletion of a *Csf1r* enhancer leads to the absence of microglia in the brain^42^ (Fig. 3a). There was an increase in the density of overall CC3^+^ cells in *Csf1r*^ΔFIRE/ΔFIRE^ mice compared to heterozygous *Csf1r*^+/ΔFIRE^ littermates (Fig. 3b,c), suggesting that the absence of microglia delayed corpse clearance. However, there was no difference in the density of OPCs or differentiating oligodendrocytes (Fig. 3d), nor in the density of CC3^+^ OPCs and CC3^+^ differentiating oligodendrocytes (Fig. 3e,f). Thus, even when dying cells accumulate in the absence of microglia, dying OPCs are extremely rare.

**Figure 3.**
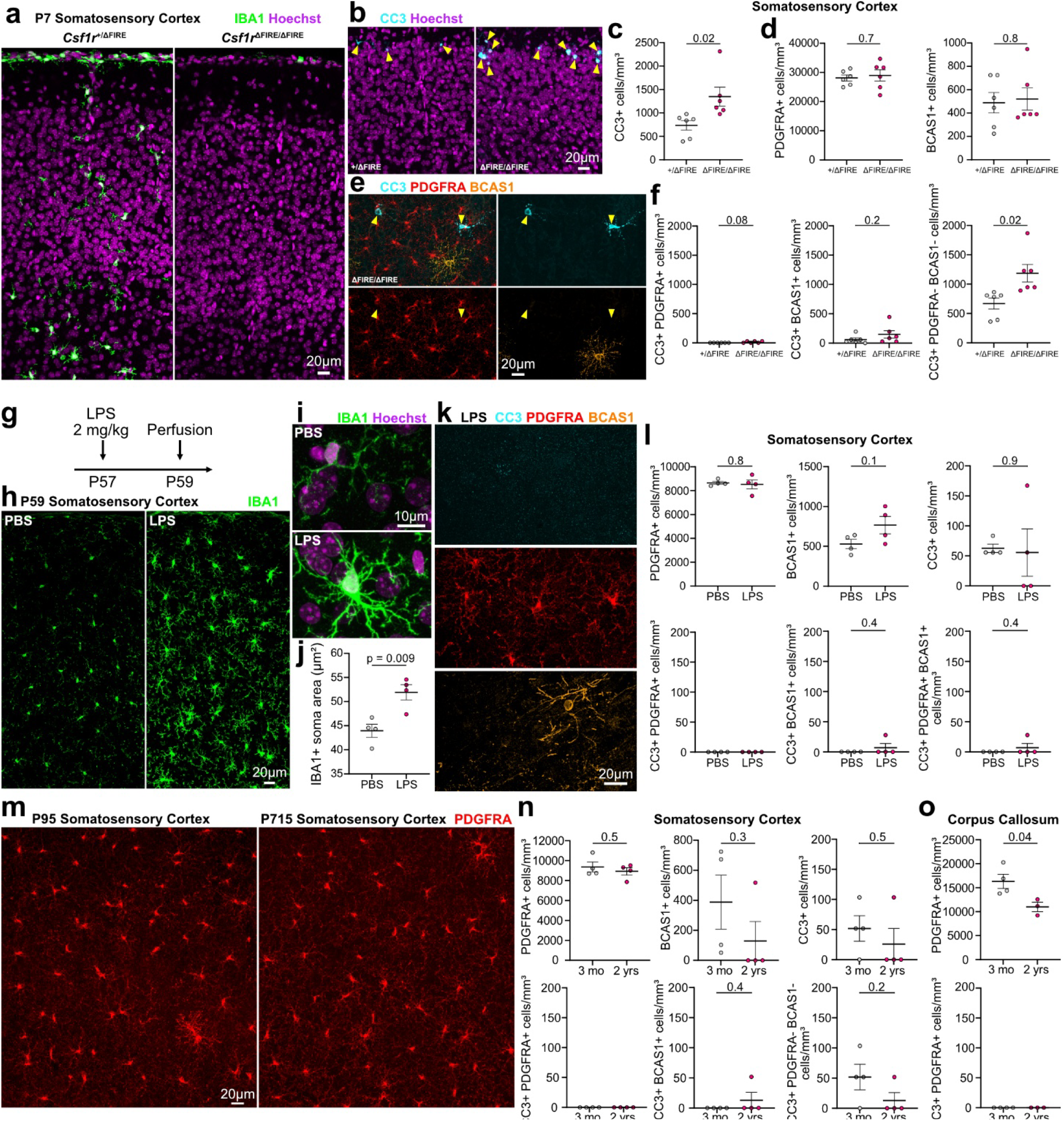
Microglia deletion, acute systemic inflammation, and aging do not reveal OPC or differentiating oligodendrocyte death. **a** Microglia (IBA1^+^; green) are absent from the brain of *Csf1r*^ΔFIRE/ΔFIRE^ mice. **b** CC3^+^ apoptotic cells (cyan; indicated with yellow arrowheads) are increased in *Csf1r*^ΔFIRE/ΔFIRE^ mice. **c** Density of CC3^+^ cells in heterozygous *Csf1r*^+/ΔFIRE^ and homozygous *Csf1r*^ΔFIRE/ΔFIRE^ mice (n=6 mice, unpaired two-tailed t-test). **d** Density of PDGFRA^+^ OPCs and BCAS1^+^ differentiating oligodendrocytes in heterozygous *Csf1r*^+/ΔFIRE^ and homozygous *Csf1r*^ΔFIRE/ΔFIRE^ mice (n=6 mice, unpaired two-tailed t-test). **e** CC3 (cyan) does not overlap with PDGFRA (red) or BCAS1 (orange) in *Csf1r*^ΔFIRE/ΔFIRE^ mice. **f** Density of CC3^+^PDGFRA^+^ cells, CC3^+^BCAS1^+^ cells, and CC3^+^PDGFRA^-^BCA1^-^ cells in heterozygous *Csf1r*^+/ΔFIRE^ and homozygous *Csf1r*^ΔFIRE/ΔFIRE^ mice (n=6 mice, unpaired two-tailed t-test). **g** Acute inflammation was induced by injecting mice with 2 mg/kg LPS. The mice were perfused 48 hours later. **h-i** Microglia morphology after a PBS or an LPS injection. **j** Microglia soma area in PBS-treated or LPS-treated mice (n=4 mice, unpaired two-tailed t-test). **k** CC3 (cyan) does not overlap with PDGFRA (red) or BCAS1 (orange) in LPS-treated mice. **l** PDGFRA^+^, BCAS1^+^, and CC3^+^ cell densities, as well as CC3^+^PDGFRA^+^, CC3^+^BCAS1^+^, and CC3^+^PDGFRA^+^BCAS1^+^ cell densities in PBS-treated and LPS-treated mice (n=4 mice; unpaired two-tailed t-tests). **m** PDGFRA^+^ OPCs in 3-month (left) and 2-year-old (right) mice. **n** PDGFRA^+^, BCAS1^+^, CC3^+^, CC3^+^PDGFRA^+^, CC3^+^BCAS1^+^, and CC3^+^PDGFRA^-^BCAS1^-^ cell densities in 3-month and 2-year-old mice (n=4 mice; unpaired two-tailed t-tests). **o** PDGFRA^+^ and CC3^+^PDGFRA^+^ cell densities in 3-month and 2-year-old mice (n=3-4 mice; unpaired two-tailed t-tests). Data shown as mean ± SEM.

Our data suggest that OPCs rarely die during development, adulthood, or remyelination. Moreover, OPCs are resistant to some toxins or stressors that induce differentiating and/or mature oligodendrocyte death, including cuprizone^26,43^ and hypoxia-ischemia^44^. Nonetheless, OPCs can die in response to injury. For example, inducing DNA damage with 2Phatal^26,39,45^ reliably leads to OPC death. In addition, treating OPC cultures or OPC-microglia co-cultures with lipopolysaccharide (LPS), or adding LPS-treated microglia conditioned medium to OPC cultures causes OPC death within 24 to 48 hours^46–51^. We therefore tested if OPCs die 48 hours after LPS-induced acute systemic inflammation in vivo (Fig. 3g). Microglia soma area was increased in the somatosensory cortex of LPS-treated mice, indicating that LPS administration induced inflammation and microglial activation^52^ (Fig. 3h-j). However, neither OPC density nor differentiating oligodendrocyte density were altered in LPS-treated mice. Moreover, we did not detect any CC3^+^ OPCs (Fig. 3k-l) nor microglia-engulfed OPCs (data not shown) in either PBS-treated or LPS-treated mice. Similarly, although we detected CC3^+^ differentiating oligodendrocytes, there was no difference between PBS-treated and LPS-treated mice (Fig. 3k-l). Taken together, our data indicate that in contrast to in vitro assays, acute systemic inflammation does not induce OPC nor differentiating oligodendrocyte death in vivo.

Aging is associated with low-grade chronic inflammation and the accumulation of DNA damage^53^. Moreover, aged OPCs express senescence markers, immune response associated genes, and cell death pathways^54–57^. We therefore wondered if, like mature oligodendrocytes^4,14^, OPCs die in aging. In line with previous studies^5,40^, OPC density did not differ in the somatosensory cortex of three-month-old and two-year-old mice (Fig. 3m-n). Moreover, we did not detect any CC3^+^ OPCs in the cortex of either three-month-old or two-year-old mice (Fig. 3n). While there were some CC3^+^ differentiating oligodendrocytes in the somatosensory cortex of two-year-old mice, their density did not differ from three-month-old mice (Fig. 3n), consistent with the low levels of differentiating oligodendrocyte death throughout life (Extended Data Fig. 1). OPC density was reduced in the aged corpus callosum, consistent with previous work^40^, but there was no evidence of CC3^+^ OPCs in either three-month-old or two-year-old mice (Fig. 3o). In the absence of OPC death, this reduced density could be due to decreased OPC proliferation in aged mice^55^ combined with the increase in callosal volume with aging^58^. Overall, our data suggest that even in aging, OPC death is extremely rare.

We found that OPCs almost never die during development, adulthood, remyelination, acute systemic inflammation, or aging. Thus, OPC homeostasis does not depend on programmed cell death. Instead, OPC density may be determined by a combination of tissue size, migration, proliferation and differentiation, which all change with development^3,16,21,55,58–65^. In line with this, global knockouts of proapoptotic genes increase the absolute number of both neurons and OPCs, but maintain the axon-to-OPC ratio, suggesting that axonal cues rather than OPC death could be a stronger regulator of OPC population homeostasis^66^. Our data also highlight that differentiating oligodendrocytes are the only stage in the oligodendrocyte lineage to face a survival decision. This is consistent with recent evidence suggesting that this fate decision is an important determinant of myelination and myelin repair^15,16,18^. Overall, our findings indicate that OPCs do not undergo spontaneous programmed cell death and challenges this assumption built into our understanding of OPC and oligodendrocyte dynamics.

## Acknowledgements

We thank members of the Hill and Hoppa labs at Dartmouth for helpful discussions and feedback on the project, and Harshini Sivagnanam for sharing the soma segmentation and area measurement Fiji macro. This work was supported by the National Institutes of Health R01NS122800 and R01NS140248 and the Esther A. & Joseph Klingenstein Fund and Simons Foundation to R.A.H. and a Postdoctoral Fellowship FG-2307-42173 from the National Multiple Sclerosis Society to Y.K.

## Author Contributions

Y.K. and R.A.H. conceived and designed the study and secured funding. Y.K. performed most of the experiments and analysis. A.S. contributed to the quantification in Fig. 2d, the staining and imaging for Extended Data Fig. 1e,f and established the TUNEL protocol. Y.K. and R.A.H. wrote the paper and all authors reviewed the manuscript.

## Data and materials availability

All data are available upon request.

## Methods Animals

All procedures were approved by the Institutional Animal Care and Use Committee at Dartmouth College (protocol # 00002158). Both male and female mice were used. When applicable, animals of both sexes were assigned to each experimental group based on genotype and availability. Data from both sexes were analyzed together and sex differences were not formally tested. Intravital imaging experiments were performed in *Cnp*-mEGFP mice^67^ (JAX: 026105) crossed with NG2-CreER™ mice^60^ (JAX: 008538) and Ai9 mice^68^ (JAX: 007909). A single 50 µL injection of 20 mg/mL tamoxifen was administered between P24 and P26 to induce recombination and imaging began between P51 and P70. The initial characterization of OPC and differentiating oligodendrocyte death in fixed tissue was performed in various strains in the colony without expected phenotypes (PV^Cre^ mice^69^ (JAX: 008069), C57BL/6J mice (JAX: 000664), and *Cnp*-mEGFP : NG2-CreER™ : Ai9 mice). Mice were perfused-fixed at P4, P7, P10, P23, P32, three months (P90-P99), one year (P366-P378) and ∼two years (P661-798). The microglia deletion experiment was performed with *Csf1r*^ΔFIRE^ mice^42^ (JAX: 032783) and the acute systemic inflammation experiment was performed in *Cnp*-mEGFP : Ai9 mice. Food and water were provided ad libitum, and mice were housed in a temperature (22 ºC) and humidity (30-70% relative humidity) controlled animal vivarium.

### Surgical Procedures

Cranial window surgeries were performed as previously described^70^. Briefly, mice were anesthetized with an intraperitoneal injection of 100 mg/kg ketamine and 10 mg/kg xylazine. 5 mg/kg carprofen was administered subcutaneously at the start and the end of surgery, and at 24 and 48 hours after the surgery. The skin covering the skull was shaved, sterilized, and then removed. A high-speed drill was used to perform a 3 mm-diameter craniotomy. The dura was removed and the skull was replaced with a no. 0 coverslip. A nut was affixed to the skull with cyanoacrylate glue and dental cement to head-fix the mice during daily intravital imaging.

### Intravital Imaging

Mice were anesthetized with 2% isoflurane and anesthesia was maintained with 1.3-1.8% isoflurane. The mice were head-fixed during imaging. Daily imaging was performed on a two-photon microscope (Bruker with Prairie software v5.4) equipped with an InSight X3 femtosecond pulsed laser (Spectra Physics) and a 20X water immersion objective (NA 1.0, Zeiss). mEGFP was excited with 920 nm and tdTomato was excited with 1040 nm. Emissions were collected with two GaAsP detectors and images were acquired at 512 pixels x 512 pixels. mEGFP and tdTomato were acquired sequentially. 345 µm to 360 µm z-stacks starting in cortical layer I were acquired with a 1.5 µm-step size. The top 150 µm were used for OPC fate and soma morphology analysis.

### Acute Systemic Inflammation

P57 mice were injected with 2 mg/kg lipopolysaccharide^71^ (LPS; Sigma, L2880) or PBS and perfused-fixed 48 hours later at P59.

### Immunohistochemistry

Mice were anesthetized with 100 mg/kg ketamine and 10 mg/kg xylazine before intracardiac perfusion with ice-cold 4% paraformaldehyde (PFA) in PBS. Neonatal mice (P4-P10) were deeply anesthetized with isoflurane or hypothermia and either perfused-fixed as above, or their brain was immediately dissected and drop-fixed in 4% PFA overnight at 4 °C. Optic nerves were dissected from neonatal mice following intracardiac perfusion with 4% PFA. Dissected brains and optic nerves were postfixed in 4% PFA overnight at 4 °C. 75 µm-thick coronal brain sections were cut with a vibrating microtome (Leica VT1000S) and stored at -20 °C in cryostorage solution (1% polyvinylpyrrolidone, 30% sucrose, and 30% ethylene glycol in 0.2 M sodium phosphate buffer, pH 7.4). Prior to staining, the sections were washed three times in PBS, for 10 minutes each. Then, the coronal brain sections or whole-mount optic nerves were incubated overnight at room temperature in primary antibodies diluted in 1% bovine serum albumin (BSA) and 0.3% Triton-X 100 in PBS. Next, the coronal brain sections or whole-mount optic nerves were washed three times in PBS, for 10 minutes each, and incubated for one (sections) or two (optic nerves) hours at room temperature in secondary antibodies diluted in 1% BSA and 0.3% Triton-X 100 in PBS. Following two 10-minute washes in PBS, the sections and optic nerves were incubated in Hoechst 33342 (Thermo Fisher Scientific, H3570, 1:2000) diluted in PBS for 20 minutes at room temperature. Following a final 10-minute wash in PBS, the sections and optic nerves were mounted on glass slides using ProLong Diamond Antifade Mountant (Thermo Fisher Scientific, P36970). All washes and incubation steps were performed on a rotating shaker.

The following primary antibodies were used: goat anti-PDGFRA (R&D Systems, AF1062, 1:1000), rabbit anti cleaved caspase-3 (Cell Signaling Technology, 9661S, 1:700), rabbit anti Iba1 (Wako, 019-19741, 1:1000), guinea pig anti-BCAS1 (Synaptic Systems, 445 004, 1:750), guinea pig anti-CNP1 (Synaptic Systems, 355 004, 1:500), guinea pig anti-SOX10 (Synaptic Systems, 347 404, 1:500). The following secondary antibodies were used at 1:500: Alexa Fluor® 488 AffiniPure® Donkey Anti-Guinea Pig IgG (H+L) (Jackson ImmunoResearch, 706-545-148), Donkey anti-Goat IgG (H+L) Cross-Adsorbed Secondary Antibody, Alexa Fluor™ 488 (ThermoFisher, A11055), Alexa Fluor® 488 AffiniPure® Goat Anti-Guinea Pig IgG (H+L) (Jackson ImmunoResearch, 106-545-003), Donkey anti-Goat IgG (H+L) Cross-Adsorbed Secondary Antibody, Alexa Fluor™ 555 (ThermoFisher, A21432), Donkey anti-Rabbit IgG (H+L) Cross-Adsorbed Secondary Antibody, Alexa Fluor™ 555 (ThermoFisher, A21429), Goat anti-Mouse IgG (H+L) Cross-Adsorbed Secondary Antibody, Alexa Fluor™ 555 (ThermoFisher, A21422), Alexa Fluor® 647 AffiniPure® Donkey Anti-Guinea Pig IgG (H+L) (Jackson ImmunoResearch, 706-605-148), Donkey anti-Rabbit IgG (H+L) Highly Cross-Adsorbed Secondary Antibody, Alexa Fluor™ 647 (ThermoFisher, A31573), Goat anti-Rabbit IgG (H+L) Cross-Adsorbed Secondary Antibody, Alexa Fluor™ 647 (ThermoFisher, A21244).

For stainings shown and quantified in Figures 2a-d and 3, the sections were incubated in a 10 mM copper(II) sulfate and 50 mM ammonium acetate solution, pH 5, for one hour at room temperature on a rotating shaker to quench endogenous fluorophores and lipofuscin^72^. These sections also underwent antigen retrieval, where they were incubated for two minutes at 98 ºC in pre-heated 10 mM Tris, 1 mM EDTA, and 0.05% Tween 20 in PBS, pH 9 prior to primary antibody incubation.

### Terminal Deoxynucleotidyl Transferase-dUTP Nick End Labeling (TUNEL)

TUNEL was performed using a Click-It Plus TUNEL assay (Invitrogen, C10617), according to the manufacturer’s instructions. Incubation times were modified to optimize labeling in free-floating 75 µm brain sections. Briefly, TUNEL was performed following antibody labeling as described above, immediately before incubation with Hoechst 33342. Coronal brain sections were washed three times in PBS for 10 minutes each. Next, the sections were placed on glass slides, surrounded by a hydrophobic barrier and stained directly on the slides. The sections were incubated in 4% PFA pre-heated to 37 ºC for 15 minutes. Then, the sections were incubated in Proteinase K for 20 minutes. Following a 5-minute wash in PBS, the sections were permeabilized in 0.3% Triton-X 100 in PBS for one hour. Following two 10-minute washes in PBS, sections were fixed for 5 minutes in 4% PFA pre-heated to 37 ºC. After a further two 5-minute washes in PBS, the sections were incubated in Terminal deoxynucleotidyl Transferase (TdT) Reaction Buffer for 10 minutes at 37 ºC, and subsequently incubated in TdT reaction mixture for one hour at 37 ºC. The sections were then washed in 3% BSA and 0.1% Triton-X 100 in PBS for 5 minutes, and incubated in Click It Plus TUNEL reaction cocktail for 30 minutes at 37 ºC. Finally, the sections were washed in 3% BSA in PBS, and following a further three 10-minute washes in PBS, incubated in Hoechst 33342 as described above.

### Fixed Tissue Imaging

Fixed tissue imaging was performed with an upright laser scanning confocal (Leica SP8 with LAS X software version 3.5.7.23225) equipped with a 20X air objective (NA 0.75, Leica) and a 63X oil immersion objective (NA 1.4, Leica). 405 nm, 488 nm, 552 nm, and 633 nm laser lines were used to excite fluorophores. Fluorophores were imaged sequentially, from longest to shortest wavelengths. Z stacks were acquired in 1.5 µm-steps when imaging with the 20X air objective and in 0.5 µm-steps when imaging with the 63X oil immersion objective. Images were acquired at 2048 pixels x 2048 pixels. Imaging fields were selected with reference to maturation stage markers only.

### OPC Fate and Soma Morphology Analysis

Time series hyperstacks were created from 21 days of daily intravital imaging in *Cnp*-mEGFP : NG2-CreER™ : Ai9 mice using Fiji^73^ (version 1.54p). z-projections were created and registered with the “Correct 3D drift” plugin^74^. The fate of tdTomato^+^ cells with OPC morphology was tracked, and four possible outcomes were noted: (1) remains an OPC; (2) differentiates as determined by *Cnp*-mEGFP onset; (3) divides; or (4) dies, as determined by soma condensation and process fragmentation followed by soma disappearance or soma disappearance alone when not initially positioned near the imaging volume edges. Dying cells and a matched number of cells that differentiated or remained OPCs were cropped out, their fate was blinded using Fiji’s File Name Encrypter, and their soma was manually traced with the brush selection tool to measure the area and circularity. These measurements were performed on the day prior to their fate. To analyze OPC fate and soma morphology during remyelination, the same fate and soma morphology analysis were performed on z-projected time series hyperstacks from a previously published dataset^14^ between days 28 and 45 after oligodendrocyte 2Phatal.

### Cell Death Staining Analysis

OPC, differentiating oligodendrocyte, and dying cell numbers were manually counted through fixed volume z-stacks, which were used to calculate densities in Figs. 2 and 3n-o and Extended Data Fig. 1. Six non-overlapping images from two brain slices per mouse were analyzed in the somatosensory cortex (from the pial surface to ∼500 µm below, cortical layers ∼I-IV). Two images of the corpus callosum midline from two brain slices per mouse were analyzed. The bounds of the callosal area to analyze were manually traced based on the presence of aligned nuclei and/or myelin. In Fig. 2b, apoptotic morphology was defined by a condensed soma, condensed and smooth nucleus, and fragmented processes. Whole optic nerves (one per mouse) were analyzed to measure the total CC3^+^ cell number. Four non-overlapping ROIs per optic nerve were used to quantify the number of CC3^+^ OPCs and oligodendrocyte lineage cells. In Fig. 2h, the number of engulfed OPCs was counted manually in z-projections and engulfment was verified in the z-stack. In Fig. 3a-l, CC3^+^ cells and differentiating oligodendrocytes were counted manually through fixed volume z-stacks, as above. Total OPC numbers were counted in z-projections using a custom Fiji macro and manually corrected as needed. Claude Sonnet 5.5 (Anthropic) was used to help write the Fiji macro. Images were blinded with Fiji’s File Name Encrypter so that the experimenter was blinded to the age or treatment condition in all experiments.

### Microglia Soma Area Analysis

Six non-overlapping images from two brain slices per mouse were analyzed in the somatosensory cortex (from the pial surface to ∼500 µm below) of LPS-treated or vehicle-treated mice. Five microglia were randomly selected in each image, and a z-projection of 3 slices was cropped out for each cell. The cell soma was segmented with a custom Fiji macro and manually corrected with the brush selection tool as needed for area measurements. Claude Opus 4.5 (Anthropic) was used to help write the Fiji macro. Images were blinded with Fiji’s File Name Encrypter so that the experimenter was blinded to the treatment group.

### Statistics and Reproducibility

Data are shown as mean ± standard error of the mean (SEM). Statistics were performed with GraphPad Prism (versions 10.6.0 to 11.0.2). In Fig. 1, Tukey’s multiple comparison test was used following a statistically significant one-way ANOVA. Variance was tested with a Brown-Forsythe test and did not differ between groups. In Fig. 3, unpaired two-tailed t-tests were used. Variance was tested with an F test, and Welch’s t-test was used when variance was unequal. Proportions of dying differentiating oligodendrocytes and OPCs were compared with Yates’ continuity-corrected χ^2^ test.

## Extended Data Figures

**Extended Data Figure 1.**
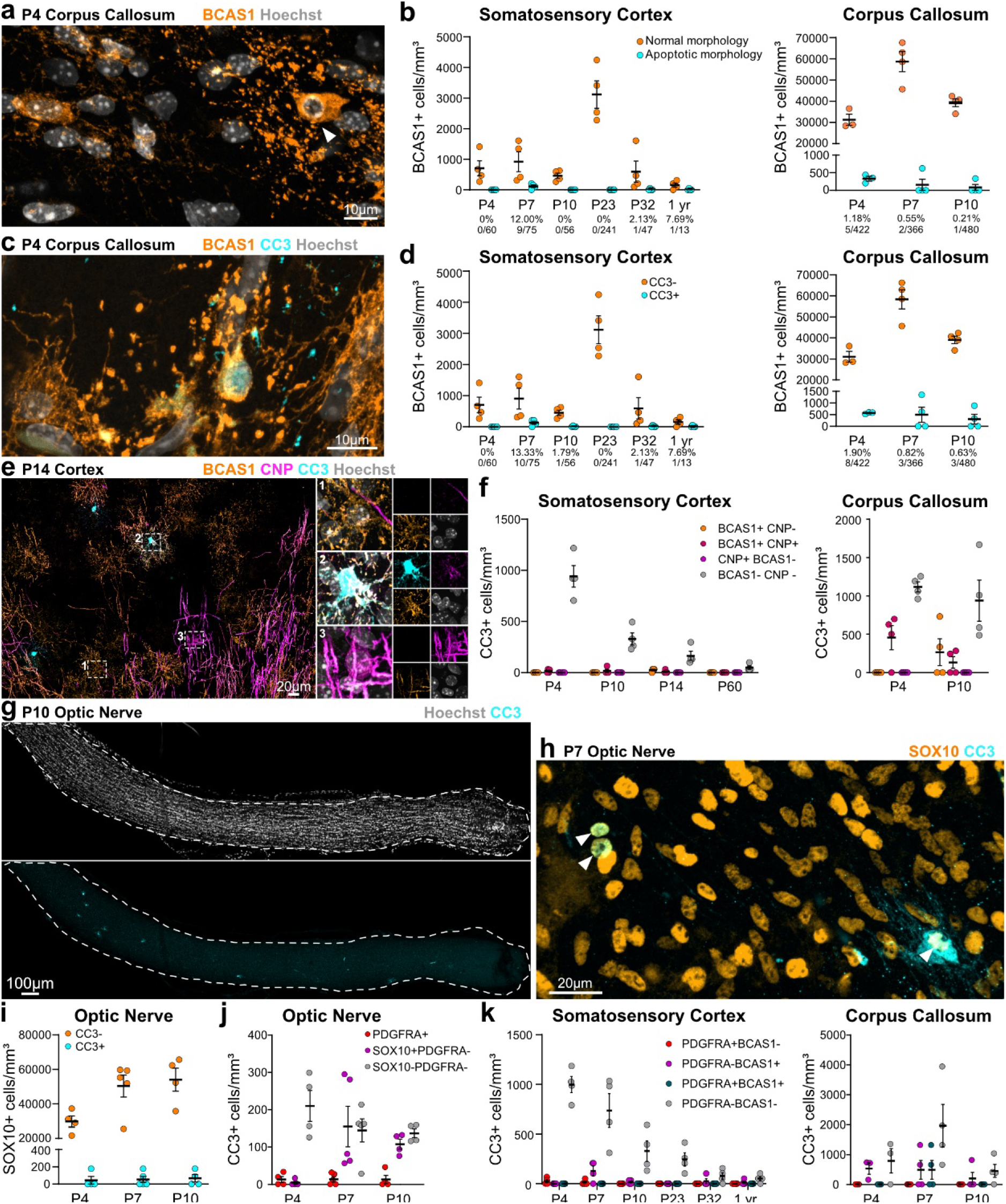
Low levels of differentiating oligodendrocyte death throughout life. **a** Example of a BCAS1^+^ differentiating oligodendrocyte with a condensed nucleus and fragmented processes (white arrowhead) and a BCAS1^+^ differentiating oligodendrocyte with a normal morphology in the P4 corpus callosum. **b** Density of BCAS1^+^ differentiating oligodendrocytes with a normal (orange), or apoptotic morphology (cyan) (n=3-4 mice). The numbers below the x axis indicate the proportion of BCAS1^+^ cells with an apoptotic morphology/BCAS1^+^ cells. **c** Example of a BCAS1^+^CC3^+^ cell in the P4 corpus callosum. **d** Density of BCAS1^+^CC3^+^ (orange) and BCAS1^+^CC3^-^ (cyan) cells (n=3-4 mice). The numbers below the x axis indicate the proportion of BCAS1^+^CC3^+^ cells/BCAS1^+^ cells. **e** Example BCAS1, CNP, and CC3 staining. Boxed cell 1 is a BCAS1^+^CNP^-^CC3^-^ differentiating oligodendrocyte. Boxed cell 2 is a BCAS1^+^CNP^+^CC3^+^ differentiating oligodendrocyte. Boxed cell 3 is a BCAS^-^CNP^+^CC3^-^ oligodendrocyte. **f** Identity of CC3^+^ cells (n=4 mice). **g** Single z-slice from a whole-mount P10 optic nerve with an example of CC3 staining. **h** Example SOX10 and CC3 staining. The white arrowheads indicate SOX10^+^CC3^+^ cells. Overall, there is little overlap between SOX10 and CC3. **i** Density of SOX10^+^CC3^-^ (orange) and SOX10^+^CC3^+^ (cyan) cells (n=4-5 mice). The numbers below the x axis indicate the proportion of SOX10^+^CC3^+^ cells/SOX10^+^ cells. **j** Identity of CC3^+^ cells (n=4-5 mice). **k** Identity of CC3^+^ cells (n=3-4 mice). All data are shown as mean ± SEM.

**Extended Data Figure 2.**
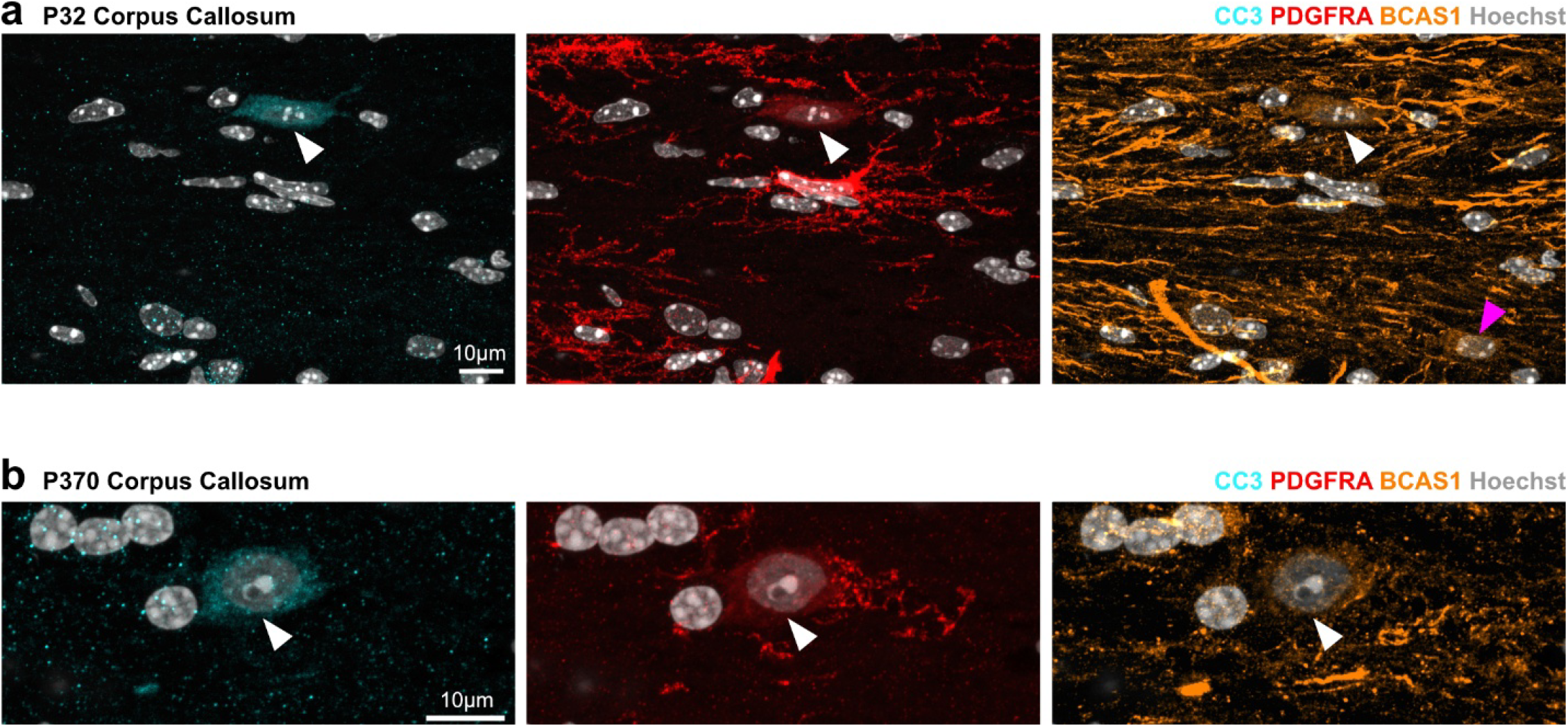
Dying OPCs in the adult corpus callosum are BCAS1 positive. **a** The white arrowhead indicates the sole PDGFRA^+^CC3^+^ cell detected in the P32 corpus callosum. The cell is also BCAS1^+^, indicating that it may have started to differentiate. The magenta arrowhead indicates an example BCAS1^+^PDGFRA^-^CC3^-^ cell. **b** The white arrowhead indicates the sole PDGFRA^+^CC3^+^ cell detected in the one-year-old corpus callosum. The cell is also BCAS1^+^, indicating that it may have started to differentiate.

**Extended Data Figure 3.**
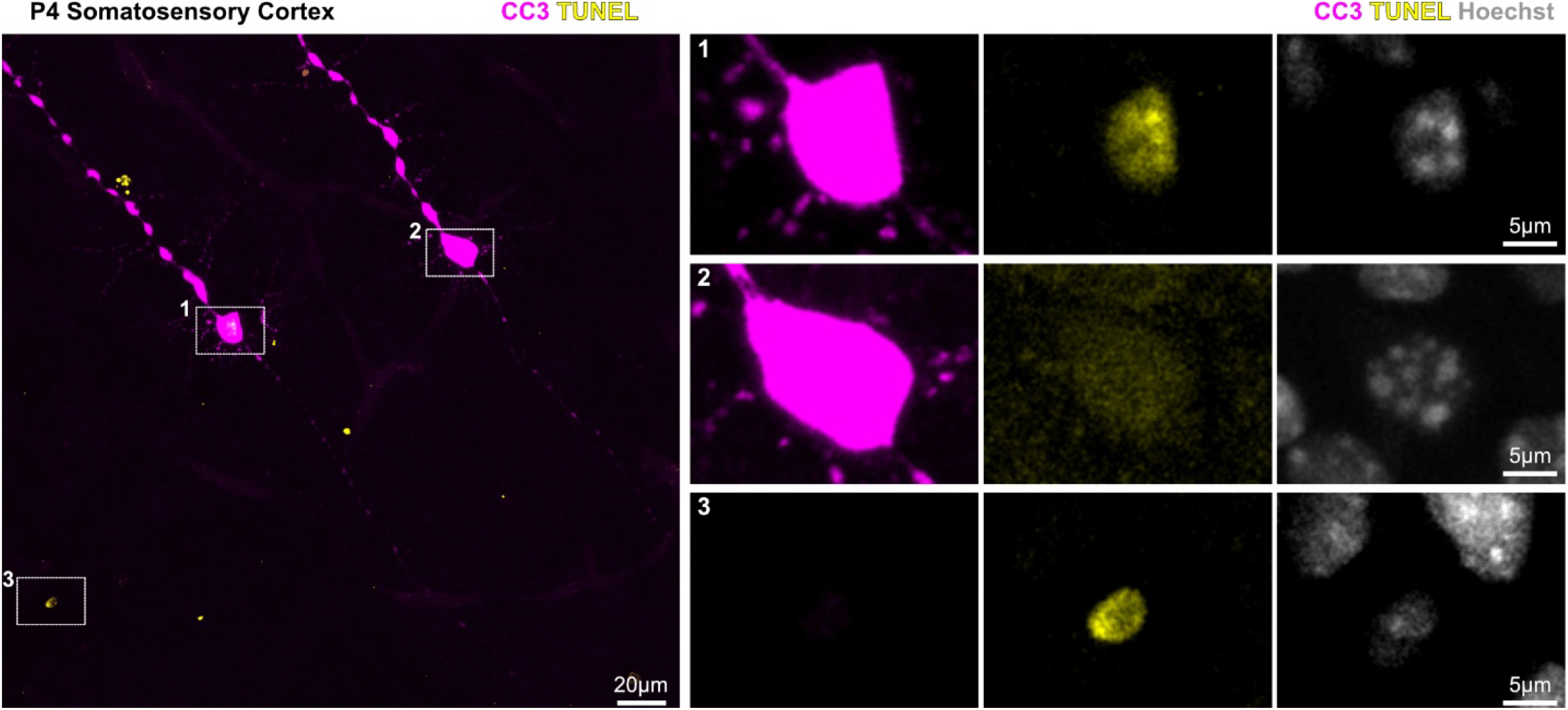
TUNEL and CC3 label different cell populations. Example of TUNEL and CC3 labeling in the P4 somatosensory cortex. CC3 labels dying neurons (boxed cells 1 and 2), but cell 1 is the only neuron that is also TUNEL^+^. Boxed cell 3 is TUNEL^+^, but CC3^-^.

